# Competition Between Public Resources Promotes Cooperation

**DOI:** 10.1101/2020.09.30.320382

**Authors:** Mohammad Salahshour

**Affiliations:** Max Planck Institute for Mathematics in the Sciences, Inselstrasse 22, D-04103, Leipzig, Germany

**Keywords:** Cooperation, Public Goods Game, Evolutionary Games

## Abstract

As cooperation incurs a cost to the cooperator for others to benefit, its evolution seems to contradict natural selection. How evolution has resolved this obstacle, has been among the most intensely studied questions in the evolutionary theory in recent decades. Here, by showing that competition between public resources provides a simple mechanism for cooperation to flourish, we uncover a novel road to the evolution of cooperation. Such a mechanism can be at work in many biological or social contexts where individuals can form different groups or join different institutions to perform a collective action task, or when they can choose between different collective actions, with different profitability. As a simple evolutionary model suggests, in such a context, defectors tend to join the highest quality resource. This allows cooperators to survive and out-compete defectors by sheltering in a lower quality resource. Cooperation level is maximized however, when the qualities of the two highest quality resources are similar, and thus, they can perform the most competitively to attract individuals.

## Introduction

The Public goods game, also known as *n* person prisoner’s dilemma, has been one of the paradigms used in many studies on the evolution of cooperation (1–14). This game is played among *n* individuals. Each individual can either cooperate or defect. Cooperators contribute an amount *c* to the public good. Defectors contribute nothing. All the contributions are multiplied by an enhancement factor *r < n* and are distributed equally among the players. When played in an evolutionary context in a well-mixed population, defectors, not paying the cost of cooperation, reach the highest payoff, and dominate the population. However, contrary to this expectation, empirical evidence suggests from bacteria to animals and humans a high level of cooperation has evolved in the course of evolution (15–18). As a result of many attempts to resolve this apparent paradox, some mechanisms which are capable of promoting cooperation in public goods game have come into the light (19–22). Examples include indirect reciprocity, such as reputation effects (1, 2), network and spatial structure (3, 4), voluntary participation (4–6), reward (7–9), and possibly punishment (10–14). As we show here, competition between public resources provides yet another road to the evolution of cooperation. This mechanism can be at work in a situation where different public resources compete to attract individuals. Such a situation can naturally occur in many examples of collective action dilemmas observed in animal and human societies (24). For instance, different public resources can be different collective actions with different profitability, different groups an individual can join to perform a collective action task, such as fishing or foraging (23, 24), or different places or groups an individual can join to perform an economic activity (25–28). As our analysis shows, in such a situation, defectors tend to join the highest quality public resource. This enables cooperators to survive and out-compete defectors by sheltering in a lower quality public resource. Cooperation level is maximized however, when the two public resources have similar qualities, and thus, can perform the most competitively. Our results thus show, being able to choose between different collective action tasks, and competition between different collective actions to attract individuals, provides a natural mechanism for solving collective action problems.

## The Model

To see how competition between public resources promotes cooperation, we consider a general case where *n* ≥ 2 competing public resources with enhancement factors *r*_1_ to *r*_*n*_ exist. For simplicity, we begin by a case where two different public resources with enhancement factors *r*_1_ and *r*_2_ exist. More precisely, we consider a population of *N* individuals, each having a strategy *s* (which can be cooperation (*C*), or defection (*D*)), and a preferred game *i* (which can be game 1 or game 2). At each time step, groups of *g* individuals are formed at random, from the population pool. Individuals in each group enter their preferred public goods game, play the game, and gather payoff according to the outcome of the game. In addition, they receive a base payoff *π*_0_, from other activities not related to the public goods game. After playing the games, individuals reproduce with a probability proportional to their payoff. The new generation replaces the old one such that the population size remains constant. Offspring inherit the strategy and the preferred game of their parents, subject to mutations. Mutations in the strategy and the preferred game occur independently, each with a probability *ν*. In the case that a mutation occurs, the value of the corresponding variable is flipped to its opposite value (e.g. *C* to *D* for the strategy and game 1 to game 2 for the preferred game).

## Results

As shown in the method section, the model can be solved analytically in terms of the replicator-mutation equation. Analysis of the model by simulations and numerical solution of the replicator dynamics reveals, depending on the parameter values and the initial conditions, the dynamics can settle in a fixed point or a limit cycle. We begin by studying cyclic behavior. In Fig. (1.a), we plot the density of cooperators (solid blue) and defectors (dash-dotted red) in game *i*, denoted as, respectively, 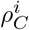 and 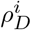, for *i* = 1 and *i* = 2, as a function of time. Here, the initial condition is a random initial condition in which both the strategy and the preferred game of the individuals are assigned at random. In Fig. (1.b) the total density of cooperators 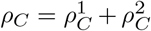 (solid blue), and defectors 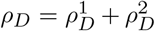 (dash-dotted red) are plotted. Here, *g* = 10, *r*_1_ = 4.4, *r*_2_ = 2.8, and *ν* = 10^−3^. Here, and in the following, we fix *c* = *π*_0_ = 1. The top panels show numerical solutions of the replicator dynamics, and the bottom panels show the results of a simulation in a population of size *N* = 10000. Comparison shows a high level of agreement between the result of simulations in a finite population and solutions of replicator dynamics, which is an exact solution of the model in the infinite population limit. Both simulation and analytical solution show that, as a result of competition between public goods to attract individuals, cooperators are preserved and cyclically dominate the population. When the density of cooperators in a public resource *i* increases, defectors in game *i* obtain a high payoff and thus increase in frequency. This decreases the payoff of cooperators in game *i* and consequently they obtain a higher payoff by switching to the other public resource. This, in turn, decreases the payoff of defectors in game *i*, and thus 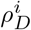 starts to decrease when 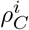 decreases enough.

**Fig. 1.**
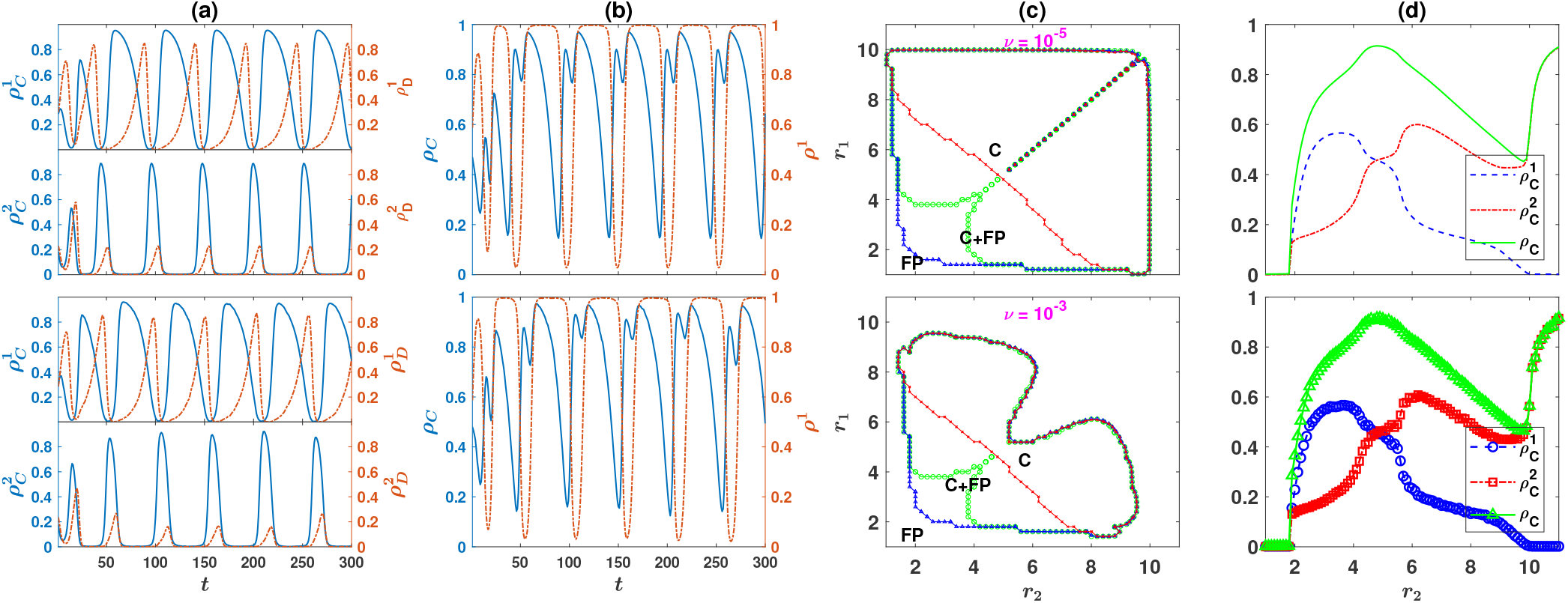
Competition between public resources promotes cooperation. (a): The density of cooperators (solid blue), 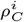, and defector (dash-dotted red), 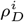, in a game *i*, for *i* = 1 and 2, as a function of time. The two top (bottom) panels result from the replicator dynamics (simulation). (b): The total density of cooperators, 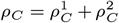 (solid blue), and defectors 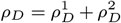 (dash-dotted red), as a function of time. The top panel results from replicator dynamics and the bottom panel results from a simulation. In (a) and (b), *g* = 10, *ν* = 10 ^*−*3^, *r*_1_ = 4.4, and *r*_2_ = 2.8. Simulations are performed in a population of size *N* = 10000. (c): The phase diagram of the model for two different mutation rates and *g* = 10, as it results from replicator dynamics. The phase diagram shows two distinct mono-stable phases where the dynamics settle into a fixed point (denoted by FP) or a limit cycle (denoted by C). In between, for small enhancement factors, the system posses a bi-stable region where both the limit cycle and the defective fixed point are stable (the region between blue (triangles) and red (crosses) lines, denoted by C+FP). The phase boundary (green line) can be defined as the line where the transition between the two phases occurs starting from a random assignment of strategies (i.e. 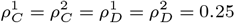). (d): The time average density of cooperators in game 1 (dashed blue), 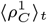, in game 2 (dash-dotted red), 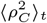, and the total density of cooperators (solid green), 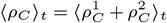, as a function of *r*_2_. The level of cooperation decreases with increasing the difference between the quality of the two resources and is maximized for *r*_2_ = *r*_1_ (in the region *r*_*2*_ < *g*). Here, *g* = 10, *ν* = 10^*−*3^, *r*_1_ = 4.8. Top panel results from replicator dynamic, and the bottom panel result from a simulation in a population of size *N* = 20000.

As mentioned before, cyclic behavior is not the only attractor of the dynamics. To see this, in Fig. (1.c), the phase diagrams of the system in the *r*_1_ − *r*_2_ plane, for *g* = 10, and two different mutation rates (indicated in the figure) are plotted. For too small enhancement factors, competition between games can not alleviate the strong disadvantage cooperators experience due to a small enhancement factor. Consequently, the dynamics settle into a (defective) fixed point in which defectors dominate the population. As either of the enhancement factors increases beyond a threshold (denoted by blue triangles), competition between games becomes strong enough to promote cyclic cooperation. In this region, the dynamics become bistable, and depending on the initial condition, the dynamics settle either on a defective fixed point or a limit cycle. Further increasing the enhancement factors, at a second bifurcation line (denoted by red crosses) the defective fixed point loses stability. The dynamics become mono-stable and settle in a limit cycle starting from all the initial conditions. The phase boundary between the defective and cyclic phases in the *r*_1_ − *r*_2_ plane can be defined as the line where the transition between the two phases occurs starting from a homogeneous initial condition (i.e. 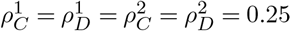). This is denoted by green circles in the figure.

By further increasing the enhancement factors, beyond a final transition line (the line appeared to be black) the limit cycle becomes unstable and the dynamics settle in a fixed point in which cooperators dominate. We note that in contrast to the transition from the defective fixed point to the cyclic orbit, the transition between the cyclic orbit and the cooperative fixed point shows no bi-stability and happens continuously (in the sense that the amplitude of fluctuations decreases gradually by increasing the enhancement factors until fluctuations vanish at the transition line). Consequently, this transition line occurs at the same value of *r*_1_ and *r*_2_ for all initial conditions. By comparison of the phase diagram for different mutation rates, it is seen that the same scenario happens for all the mutation rates. Quantitatively, however, larger mutation rates decrease the size of the region in the *r*_1_ − *r*_2_ plane where cyclic behavior can occur.

We have seen that competition between public resources promotes cooperation. An interesting question is how the level of cooperation depends on the enhancement factors? To see this, in Fig. (1.d) we fix *r*_1_ = 4.8 and plot the time average density of cooperators in game 1, 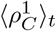 (dashed blue line), in game 2, 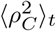 (dash-dotted red line), and the time average total density of cooperators 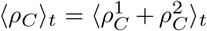 (solid green) as a function of *r*_2_. Here, *g* =10, *ν* =10^−3^, and a homogeneous initial condition is used (i.e. 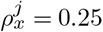 for all *j* and *x*). The top panel results from the replicator dynamics, and the bottom panel shows the result of a simulation in a population of size *N* =20000. As argued before, for small *r*_2_ the dynamics settle in a defective fixed point. By increasing *r*_2_ beyond a certain value of *r*_2_, cooperators can survive and cyclically dominate the population. The density of cooperators increases by increasing *r*_2_ and reaches its maximum, at *r*_2_ = *r*_1_ in a resonance-like phenomenon. Interestingly, increasing *r*_2_ beyond *r*_1_ has a detrimental effect on the level of cooperation. This shows the importance of competitiveness of the two alternative public resources for the evolution of cooperation, as the level of cooperation decreases with increasing the difference between the quality of the two public resources. This can be seen to be valid for all values of *r*_1_, in Fig. (2.a), where the time average of the total density of cooperators in the *r*_1_ − *r*_2_ plane, is plotted. Here and in other parts of Fig. (2), *g* = 10, *ν* =10^−3^, and a homogeneous initial condition is used. The top panels result from the replicator dynamics, and the bottom panels show the result of a simulation in a population of size *N* = 20000. As can be seen, the density of cooperators is the highest on the diagonal where *r*_1_ = *r*_2_, and decreases with increasing the distance from the diagonal (i.e. with increasing the difference of the enhancement factors of the two alternative public resources).

**Fig. 2.**
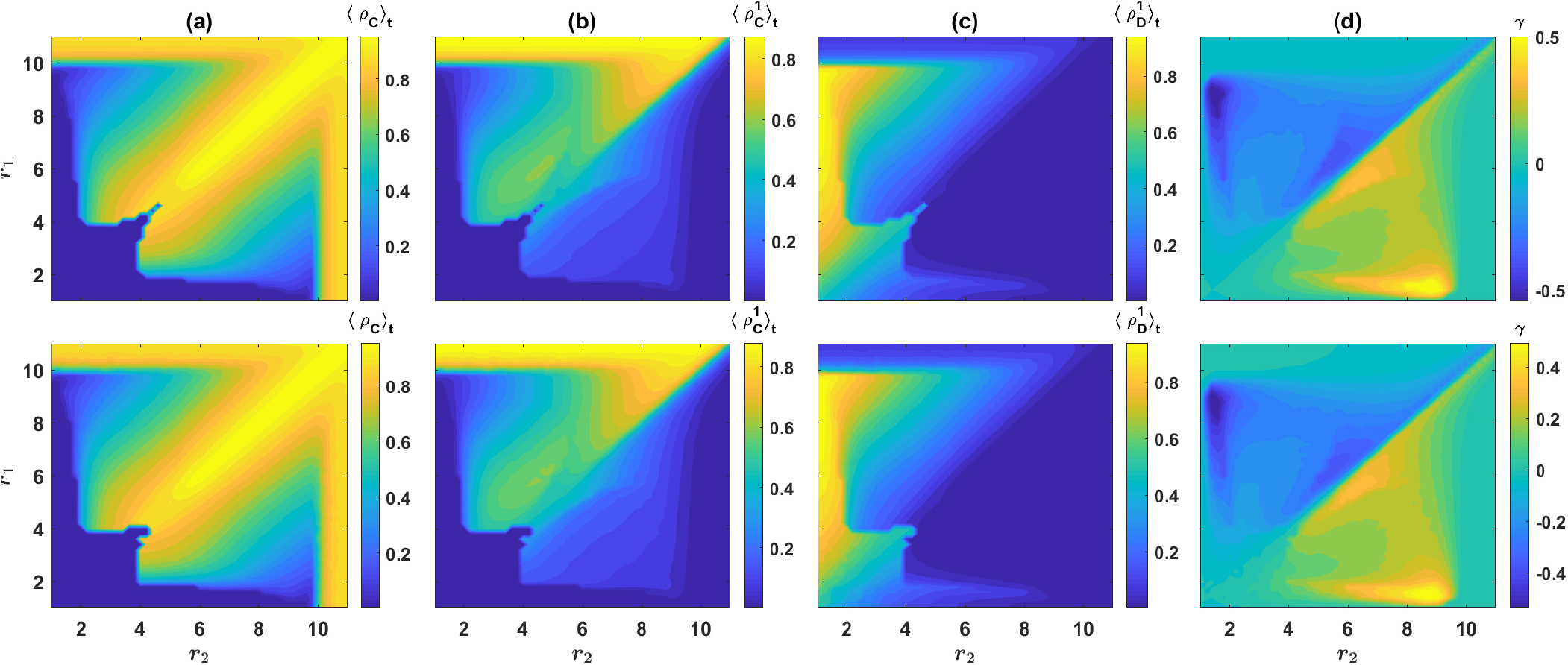
(a): The contour plot of the time average density of cooperators, 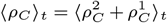. Cooperation evolves for large enough enhancement factors, and its level is maximized when the enhancement factors of the two resources are equal. (b) and (c): The time average density of cooperators (b), and defectors (c), in game 1. Compared to cooperators, defectors are more likely to prefer the highest quality resource. Consequently, cooperators can use the low quality resource as a shelter when defectors dominate. (d) Contour plot of 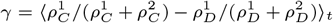. This is a measure of how much cooperator are more likely to prefer game 1, compared to defectors. Cooperators are more likely to prefer game 1, when it is the lower quality game, and defectors are more likely to prefer game 1 when it is the higher quality game. Here, *g* = 10 and *ν* = 10^−3^. Top panels result from the replicator dynamics, and bottom panels show the result of a simulation in a population of size *N* = 20000.

To shed more light on the mechanism by which competition between public resources promotes cooperation, and to see why the cooperation level is maximized when the two competing games have similar enhancement factors, in Fig. (2.b) and Fig. (2.c), we plot the time average density of, respectively, cooperators and defectors in game 1 (Due to the symmetry of the two games, the corresponding densities in game 2 result from this by reflection with respect to the diagonal). Here, it can be seen that most of the individuals prefer game 1, as long as it has a higher enhancement factor. Interestingly, the low-quality resource is often immune to invasion by defectors: while the inferior game can attract cooperators, defectors strongly prefer the higher quality game. This phenomenon can be made more quantitative by calculating the fraction of cooperators who prefer game 1 as 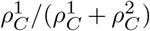, and the fraction of defectors who prefer game 1, 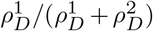. These are a measure of how much a cooperator or a defector is likely to prefer game 1. The time average difference between these two quantities, 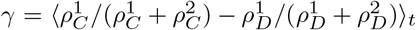 is plotted in Fig. (2.d). As can be seen, cooperators are always more likely to prefer the lower quality game, than the defectors are. Similarly, defectors are more likely to prefer the highquality game, than the cooperators are. This phenomenon is crucial for competition between resources to promote cooperation: When the density of defectors increases in the population, cooperators can shelter in the lower quality resource and make it the more profitable one, and thus, by enjoying a higher growth rate, out-compete defectors. However, when the quality of the inferior resource is much lower than that of the superior resource, it becomes more difficult for cooperators to make the inferior resource the more profitable one. This, in turn, leads to a reduction in the cooperation level, by increasing the difference between the quality of the resources.

Here, we have considered a case where *n* = 2 public resources compete. As mentioned before, the model can be extended to a general case where individuals can choose among an arbitrary number, *n*, of public resources. In the Supplemental Material, we consider the *n* = 3 case, and show having a higher number of public resources for the individuals to choose among, further facilitates the evolution of cooperation. Interestingly, with *n* = 3 it can happen that the lowest quality resource becomes obsolete: While defectors predominantly choose the highest quality resource, the second quality resource can attract cooperators. The lowest quality resource on the other hand, remains deserted when its quality is sufficiently lower than the other two resources. Nevertheless, surprisingly, the possibility to choose a third low-quality resource can facilitate the evolution of cooperation, even when individuals do not choose such a low-quality resource in practice [See Supplementary figures S.4 to S.5 in the Supplemental Material].

Finally, in the Supplemental Material we argue for the robustness of our result for other parameter values. In particular, finite size analysis of the model reveals, while competition between public resources promotes cooperation in small population sizes as well, this mechanism is stronger in large population sizes [See Supplementary figures S.2]. In this regards, competition between public resources seems to contrast many mechanisms for the evolution of cooperation which suffer from the so-called scalable cooperation problem (29, 30), according to which cooperation diminishes in large population sizes.

## Conclusion

We have seen that in a situation where individuals can choose between different public resources, competition between public resources promotes cooperation. In such a context, while defectors predominantly prefer the highest quality resource, cooperators show an inclination to choose lower quality resources as well, where they can work cooperatively to out-compete defectors. Cooperation level however, is maximized, when the two highest quality resource have similar qualities, and thus, can perform the most competitively. Our analysis thus shows, competition between public resources can fruitfully be at work to give rise to a high level of cooperation not only in small scale societies, but more prominently, in many large scale societies where individuals can choose among different groups or activities, possibly with different profitability, to engage in a collective action.

## Materials and Methods

### The replicator dynamics

The replicator-mutation dynamics for the model in the general case that individuals can choose among *n* different public resources can be written as follows:

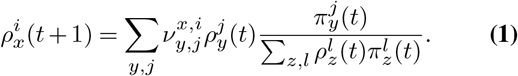

Here, *x*, *y*, and *z* refer to strategies and can be either *C* or *D*, and *i*, *j*, and *l* refer to the public resources numbered from 1 to *n*. 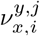 is the mutation rate from a strategy combination which prefers public resource *i* and plays strategy *x* to a strategy combination which prefers public resource *j* and plays strategy *y*. These can be written in terms of mutation rates. In the case of *n* = 2 public resources, these read as follows:

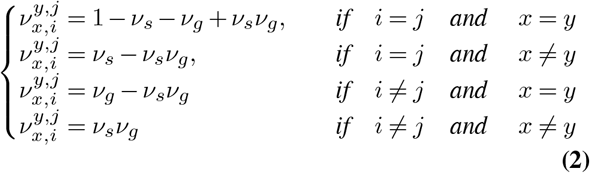

Here, we have assumed that the mutation rate in the strategies is *v*_*s*_ and the mutation rate in the game preference is *v*_*g*_ (through this study we set *ν*_*s*_ = *v*_*g*_ = *ν*). In eq. (S.1), 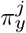 is the expected payoff of an individual who prefers public resource *j* and plays strategy *y*. These can be written as:

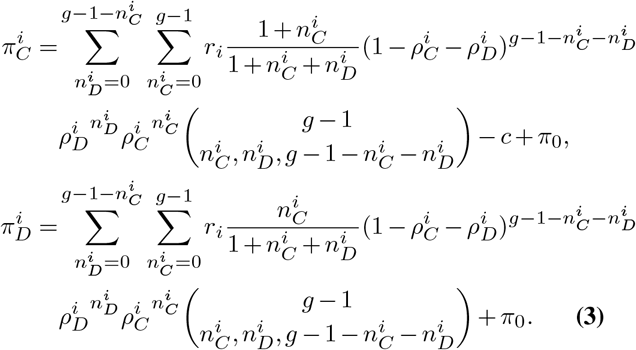

Here, 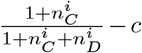 in the first equation, is the expected payoff of a cooperator who prefers the public resource *i*, in the case that 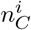 cooperators and 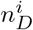 defectors in the group prefer the public resource *i*. Similarly, 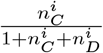 in the second equation, is the expected payoff of a defector who prefers the public resource *i*, in the case that 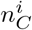 cooperators and 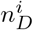 defectors in the group prefer the public resource *i*. 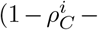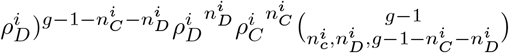 is the probability that this event occurs. 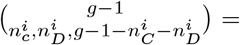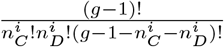 is the multinomial coefficient and is the number of ways that 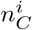 cooperators and 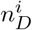 defectors who prefer game *i* can be chosen among *g* − 1 group-mates of a focal individual. Summation over all the possible configurations gives the expected payoff of a cooperator (or defector) who prefers game *i*. As all the individuals receive a base payoff *π*_0_, all the payoffs are added by this base payoff.

### The simulations and methods

Simulations used in Fig. (1.d) and Fig. (2) are run for *T* = 5000 time steps, and the time averages are taken over the last 4000 time steps. For the solutions of the replicator dynamics presented in these figures, the replicator dynamics is numerically solved for *T* = 4000 time steps, and the averages are taken in the time interval *t* = 2000 to *t* = 4000. To derive the phase diagram presented in Fig. (1.c), the following procedure is used. In order to determine the boundaries of the bistable region where both defective fixed points and cyclic behavior are possible, we note that the most favorable initial condition for the evolution of cooperation is the one in which all the individuals prefer the same game (for example (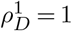, 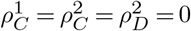). Analytical solutions starting from this initial condition give the lower boundary of the coexistence region (the blue line marked with triangles in the figure). The least favorable initial condition is the one in which all the individuals are defectors and prefer the two games in equal fractions (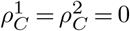 and 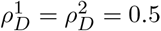). Numerical solutions of the replicator dynamics starting from this initial condition give the upper boundary of the coexistence region (the red line marked with crosses). Finally, to determine the phase boundary between the defective and cyclic phases (green line marked with circles), replicator dynamics with a homogeneous initial condition 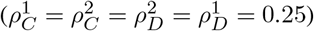) is solved. All the three sets are solved for *T* = 4000 time steps, and the values of *r*_1_ and *r*_2_ where a limit cycle appears or disappears mark the phase boundaries. As the phase boundary between the cyclic and cooperative phases for large enhancement factors shows no bi-stability, solutions starting from all the initial conditions result in the same line which appears to be black in the figure.

## ACKNOWLEDGEMENTS

The author acknowledges funding from Alexander von Humboldt Foundation in the framework of the Sofja Kovalevskaja Award endowed by the German Federal Ministry of Education and Research.

## Supplemental Material

### Supplementary Note S.1: Overview of the model

In the general case, we consider a population of *N* individuals who can play one of *n* possible public goods games (PGGs). Each individual has a preferred game *i*, which can be one of the *n* possible PGGs, and a strategy *x*, which can be cooperation (C), or defection (D). At each time step, groups of *g* individuals are drawn at random from the population pool. Individuals in each group enter their preferred PGG and play the game. That is, in each PGG, cooperators pay a cost *c* to invest an amount *c* to the public resource, and defectors pay no cost and do not invest. All the investments are multiplied by an enhancement factor (*r_i_* for PGG *i*) and are divided equally among all the individuals who participated in that PGG. In addition to the payoff received from playing the PGG, we assume individuals gather a base payoff *π*_0_ from other activities not related to the PGG. After receiving the payoffs, a selection occurs in which individuals reproduce according to their payoff. In the selection stage, the whole population is updated such that the population size remains constant. More precisely, each individual in the next generation is offspring to an individual in the past generation with a probability proportional to its payoff. Offspring inherit the strategy and the preferred game of their parent, subject to mutations. Mutations in the preferred game and the strategy of the individuals happen independently, each with probability *ν*. If a mutation in the preferred game occurs, the preferred game of the offspring is set equal to a randomly chosen PGG, other than that of its parent. In the same way, if a mutation in the strategy of an individual occurs, its strategy is set equal to a strategy other than the strategy of its parent (C to D and vice versa).

### Supplementary Note S.2: Analytical approach

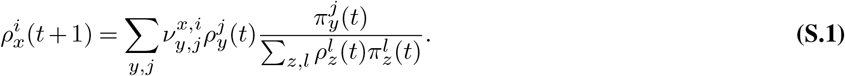

Here, *x*, *y*, and *z* refer to strategies and can be either *C* or *D*, and *i*, *j*, and *l* refer to the public resources numbered from 1 to *n*.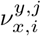 is the mutation rate from a strategy combination which prefers public resource *i* and plays strategy *x* to a strategy combination which prefers public resource *j* and plays strategy *y*. These can be written in terms of mutation rates. In the case of *n* = 2 public resources, these read as follows:

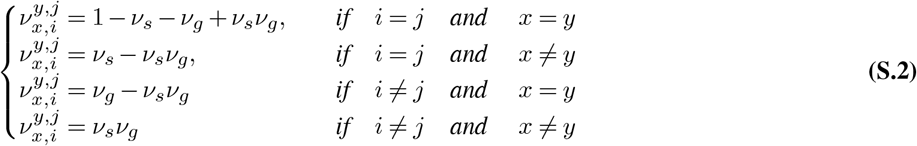

Here, we have assumed that the mutation rate in the strategies is *v*_*s*_ and the mutation rate in the game preference is *v*_*g*_ (through this study we set *v*_*s*_ = *v*_*g*_ = *v*). In eq. (S.1), 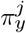 is the expected payoff of an individual who prefers public resource *j* and plays strategy *y*. These can be written as:

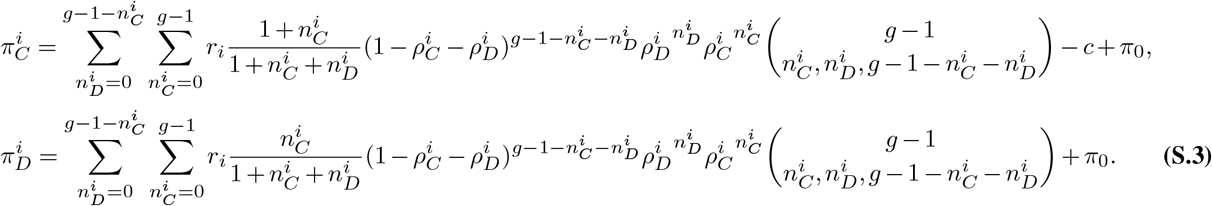

Here, 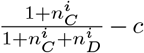 in the first equation, is the expected payoff of a cooperator who prefers the public resource *i*, in the case that 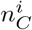 cooperators and 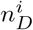 defectors in the group prefer the public resource *i*. Similarly, 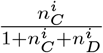, e second equation, is the expected payoff of a defector who prefers the public resource *i*, in the case that 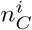 cooperators and 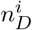 defectors in the group prefer the public resource *i*.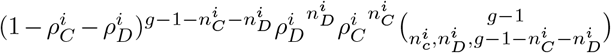 is the probability that this event occurs. 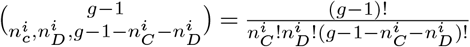 is the multinomial coefficient and is the number of ways that 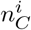 cooperators and 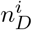 defectors who prefer game *i* can be chosen among *g* − 1 group-mates of a focal individual. Summation over all the possible configurations gives the expected payoff of a cooperator (or defector) who prefers game *i*. As all the individuals receive a base payoff *π*_0_, all the payoffs are added by this base payoff.

**Fig. S.1.**
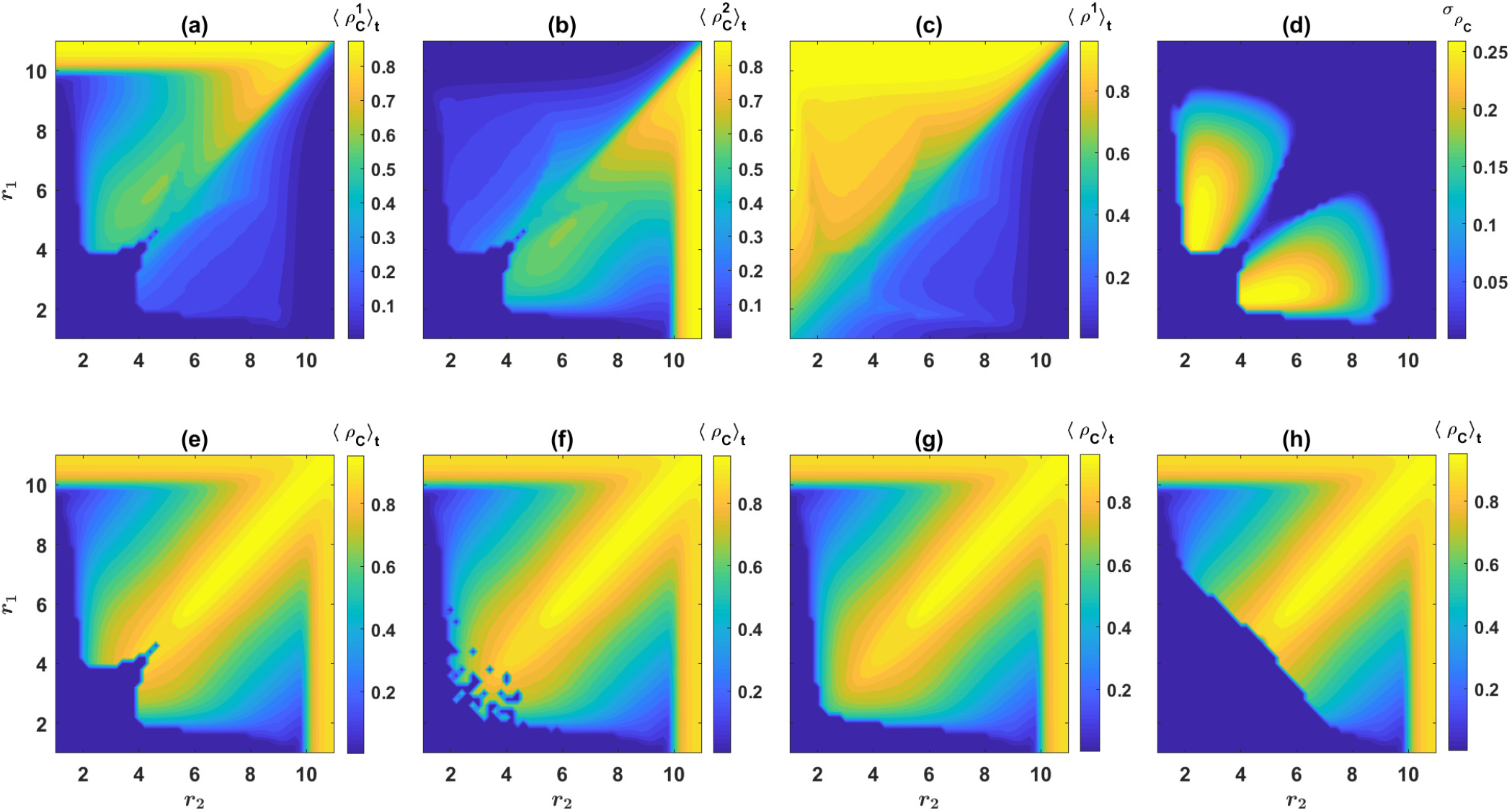
Numerical solution of the replicator dynamics. (a) to (c): (a) and (b) present color plots of respectively 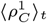 and 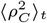, and (c) is the color plot of the time average density of individuals in game 1, 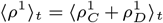. It can be seen that the density of individuals and cooperators is the highest in the higher quality resource. (d): The standard deviation of the density of cooperators, which can be used as a measure of the strength of cyclic behavior. (e) to (h): The time average density of cooperators starting from different initial conditions. In (e) the initial condition is homogeneous 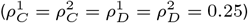, in (f), it is a randomly chosen initial condition in which 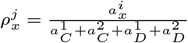, where, 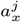 for *i* = 1 and 2, and *x* = *C* and *D*, are random numbers chosen uniformly at random in the interval [0,1]. In (g), the initial condition is a cooperation favoring one(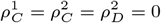 and 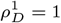), and in (h) the initial condition is a defection favoring one (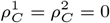 and 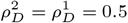). Here, *g* = 10, *ν* = 10^−3^, and *c* = *π*_0_ = 1. The replicator equations are solved for *T* = 4000 time steps and the time averages (or standard deviation) are calculated based on the last 2000 time steps. In (a) to (d), the initial condition is a homogeneous initial condition.

### Supplementary Note S.3: Simulations and derivation of the phase diagram

Simulations used in Fig. (1.d) and Fig. (2) in the main text, are run for *T* = 5000 time steps, and the time averages are taken over the last 4000 time steps. For the solutions of the replicator dynamics presented in these figures, the replicator dynamics is numerically solved for *T* = 4000 time steps, and the averages are taken in the time interval *t* = 2000 to *t* = 4000. To derive the phase diagram presented in Fig. (1.c), the following procedure is used. In order to determine the boundaries of the bistable region where both defective fixed points and cyclic behavior are possible, we note that the most favorable initial condition for the evolution of cooperation is the one in which all the individuals prefer the same game (for example 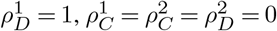). Analytical solutions starting from this initial condition give the lower boundary of the coexistence region (the blue line marked with triangles in the figure). The least favorable initial condition is the one in which all the individuals are defectors and prefer the two games in equal fractions (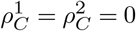 and 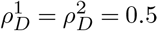). Numerical solutions of the replicator dynamics starting from this initial condition give the upper boundary of the coexistence region (the red line marked with crosses). Finally, to determine the phase boundary between the defective and cyclic phases (green line marked with circles), replicator dynamics with a homogeneous initial condition (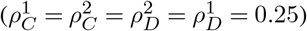) is solved. All the three sets are solved for *T* = 4000 time steps, and the values of *r*_1_ and *r*_2_ where a limit cycle appears or disappears mark the phase boundaries. As the phase boundary between the cyclic and cooperative phases for large enhancement factors shows no bi-stability, solutions starting from all the initial conditions result in the same line which appears to be black in the figure.

### Supplementary Note S.4: Analysis of the model

To perform a detailed analysis of the model with *n* = 2, we set *g* = 10, *ν* = 10^−3^, *c* = *π*_0_ = 1, and present the results of numerical solutions of the replicator dynamics in Fig. (S.1). The results of a simulation in a population of size *N* = 20000, for the same parameter values, is presented in Fig. (S.2.a) to Fig. (S.2.f). In Fig. (S.1.a) and Fig. (S.1.b), we present contour plots of, respectively, the time average cooperation level in game 1, 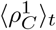, and the time average cooperation level in game 2, 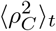. Here, and in the following, 〈·〉_*t*_ denotes a time average. In Fig. (S.1.c), the contour plot of the time average density of the individuals in game 1, 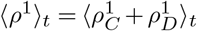 is presented. Due to the symmetry of the two games, the time average density of the individuals in game 2 is similar, and results from the reflection of Fig. (S.1.c) with respect to the diagonal. Here, the initial condition is a homogeneous initial condition in which the initial density of all the strategies equals 0.25 (that is 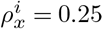 for *i* = 1 and 2, and *x* = *C* and *D*).

**Fig. S.2.**
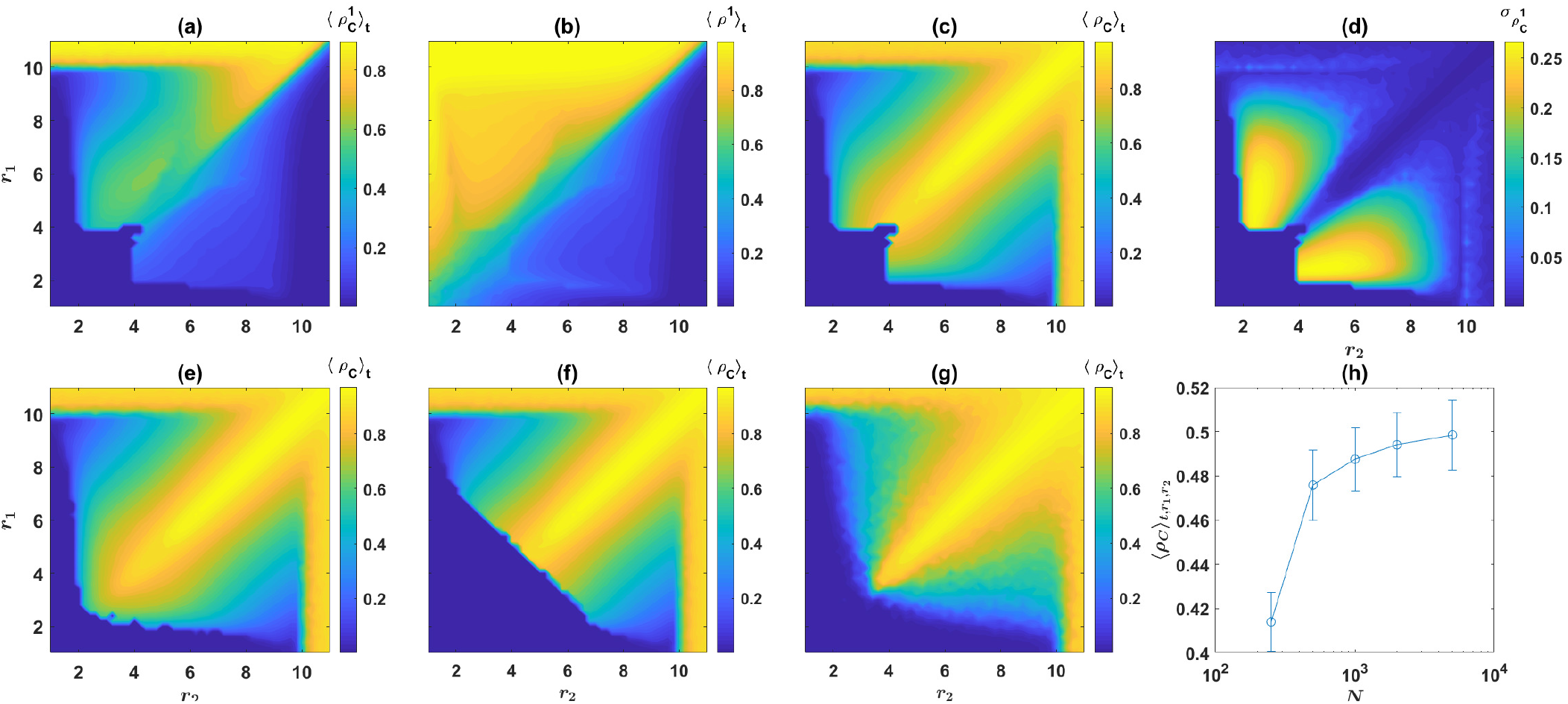
Simulations of the model. (a) and (b): (a) presents the color plot of 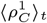 and (b) is the color plot of the time average density of individuals in game 1, 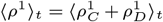. It can be seen that the density of individuals and cooperators is the highest in the higher quality resource. (c) and (d): (c) presents the time average density of cooperators starting from a homogeneous initial condition 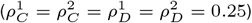), and (d) presents its standard deviation, which can be used as a measure of the strength of cyclic behavior. (e) and (f): The time average density of cooperators starting from a cooperation favoring initial condition (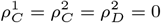 and 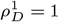) (e), and the time average density of cooperators starting from a defection favoring initial condition (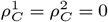and 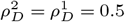) (f). (g): The density of cooperators starting from a homogeneous initial condition in a population of size *N* = 500. Comparison to (c) for *N* = 20000 shows transition to the cyclic phase occurs for smaller enhancement factors for larger population sizes. (h): The mean of the time average density of cooperators over the phase diagram (*q* ≥ *r*_1_, *r*_2_ ≤ *g*) as a function of population size. It can be seen that the level of cooperation increases with increasing the population size. Here, *g* = 10, and *ν* = 10^−3^. In (a) to (f), *N* = 20000, and the simulations are performed for *T* = 5000 time steps. In (g) *N* = 500 and the simulation is performed for *T* = 25000 steps. The averages are taken after discarding the first 500 time steps.

As can be seen in Fig. (S.1.a) and Fig. (S.1.b), the density of the cooperators is higher in the game with the higher enhancement factor. This makes the game with higher enhancement factor, on average, the more profitable one as well. As a result, as can be seen in Fig. (S.1.c), the density of individuals is higher in the game with the higher enhancement factor. Comparison with the results of simulations in Fig. (S.2.a) and Fig. (S.2.b), where respectively, the time average density of cooperators in game 1, and the density of individuals who prefer game 1, are plotted, shows the same is true for the result of simulations in finite populations. Here, as the initial condition of the simulations, strategies and the preferred games of the individuals are assigned uniformly at random (this is equivalent to a homogeneous initial condition used for the solutions of the replicator dynamics). Comparison of the results of the simulation with the solutions of the replicator dynamics shows a high agreement between the two.

To see in what parameter region the dynamics settle into a fixed point or it show cyclic behavior, we consider the standard deviation of the time series of the cooperation level in the population, 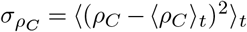. When the dynamics settle in a fixed point, this quantity equals zero. On the other hand, when the attractor of the dynamics is a cyclic orbit, *σ*_*ρC*_ takes a non-zero value. Thus, *σ*_*ρC*_ distinguishes between different attractors of the system. *σ*_*ρC*_ is plotted in Fig. (S.1.d) for the solutions of the replicator dynamics, and in Fig. (S.2.d), for a simulation with the same parameter values. Again, comparison reveals a high level of agreement between the result of simulations and the solutions of the replicator dynamics. However, we note that in the case of simulations in finite populations, due to population stochasticities, the standard deviation of the cooperation level deviates slightly from zero even in the region where the replicator dynamics settle into a fixed p oint. I n p assing, we note that, as can be seen by the sudden change in the value of *σ*_*ρC*_, while the transition from the defective fixed point to the cyclic orbit in small enhancement factors shows bistability and is discontinuous, the transition from the cyclic behavior to the cooperative fixed point in large enhancement factors shows no bistability and is gradual. That is, in large enhancement factors, by increasing the enhancement factors, the extent of fluctuations as measured by *σ*_*ρC*_ decreases continuously, until it reaches zero at the bifurcation line, and the dynamics settle into a fixed point.

**Fig. S.3.**
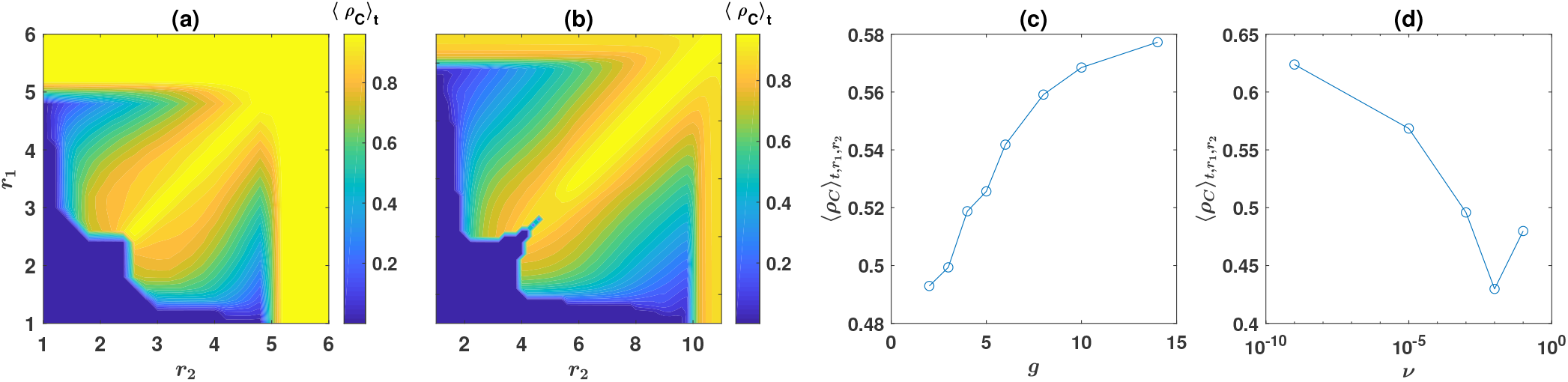
Dependence on group size and mutation rate. (a) and (b): (a) shows the time average density of cooperators 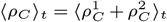 where *g* = 5 and *ν* = 10^−5^ and (b) is the case of *g* = 10 and *ν* = 10^−5^. (c): The time average density of cooperators averaged over the phase diagram (1 ≤ *r*_1_, ≤ *r*_2_ ≤ *g*) as a function of *g*. As can be seen the level of cooperation increases with *g*. Here, *ν* = 10^−5^. (d) time average density of cooperators averaged over the phase diagram (1 ≤ *r*_1_, ≤ *r*_2_ ≤ *g*) as a function of *ν*. The level of cooperation is the lowest for a comparatively large mutation rate (*ν* = 10^−2^), and increases for both smaller and larger mutation rates. Here, *g* = 10 and *π*_0_ = 1. In all the cases, the replicator equations are solved for *T* = 4000 time steps, and time averages are taken over the last *t* = 2000 time steps.

As we have seen in the main text, for small enhancement factors the dynamics show bistability and its stationary state depends on the initial condition. To see how this happens, we solve the replicator dynamics with different initial conditions and present the resulting time average cooperation level 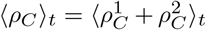, in Fig. (S.1.e) to Fig. (S.1.h). In Fig. (S.1.e), the initial condition is a homogeneous initial condition. That is, the initial density of all the four strategies are the same, and equal 0.25 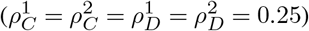. The phase boundary between the defective fixed point and the cyclic orbit can be determined based on the final state of the dynamics, starting from this initial condition. To see how the system behaves starting from different initial conditions, in Fig. (S.1.f), we present the contour plot of the time average cooperation level starting from different, randomly chosen initial conditions. Here, the *r*_1_ − *r*_2_ plane is divided into small cells, and for each cell a random initial condition is used to solve the replicator dynamics. The choice of the initial condition is such that, for each cell we set _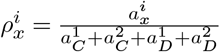_, where 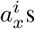 for *i* = 1 and 2, and *x* = *C* and *D*, are random numbers chosen uniformly at random in the interval [0,1]. As can be seen by comparison of Fig. (S.1.e) and Fig. (S.1.f), for both very large and very small enhancement factors, the stationary state of the dynamics does not depend on the initial condition. However, for medium enhancement factors, two different stationary states, a defective fixed point in which cooperation does not evolve, and a cyclic orbit in which cooperation evolves, are possible. Starting from a randomly chosen initial condition, the dynamics settle in one of these two attractors. Analysis of the model reveals the most cooperation favoring initial condition is an initial condition in which all the individuals prefer the same game. This can be seen by noting that, starting with such an initial condition, mutant cooperators who prefer the unoccupied game receive the highest payoff and grow in number. When the density of such cooperators increases enough, mutant defector who prefer the same game obtain the highest payoff and thus the highest growth. At this stage, the cyclic dominance of cooperators and defectors in the two games sets in. The time average cooperation level starting from one such cooperation favoring initial condition is plotted in Fig. (S.1.g). Here, the initial condition is 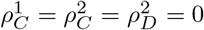 and *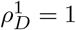*. As mentioned before, any initial condition in which all the individuals prefer the same game is equally cooperation favoring and would result in the same picture. The lower boundary of the phase diagram, that is the line above which the cyclic orbit becomes stable, can be determined starting from such an initial condition. On the other hand, the most defection favoring initial condition is the one in which all the individuals are defectors and all the different games are occupied by the same number of individuals. The time average cooperation level in the stationary state with such an initial condition is plotted in Fig. (S.1.h). This determines the upper boundary of the coexistence region above which the defective fixed point becomes unstable.

To compare the solutions of the replicator dynamics, with the results of simulations, in Fig. (S.2.c), Fig. (S.2.e), and Fig. (S.2.f), we plot the time average cooperation level, resulted from a simulation in a population of size *N* = 20000, starting from different initial conditions. In Fig. (S.2.c), the initial condition is a homogeneous initial condition, in which the strategy and the preferred games of the individuals are assigned uniformly at random, in Fig. (S.2.e), the initial condition is a cooperation favoring one. As mentioned before, this can be any initial condition in which all the individuals prefer the same game. Here, we have chosen for the initial densities of different strategies: 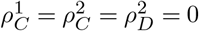 and 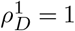. Finally, in Fig. (S.2.f), the initial condition is a defection favoring one in which we have for the initial densities of the different strategies, 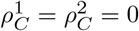and 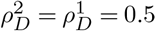. Comparison with the corresponding figures for the solutions of the replicator dynamics in Fig. (S.1), reveals a high level of agreement between the two. However, as expected, some small shifts in the positions of the phase transitions due to finite size effects are observable, especially in smaller population sizes. We return to this in the next section.

### Supplementary Note S.5: Dependence of the cooperation level on the model parameters

An important question is how the cooperation level depends on the parameters of the model? We begin answering this question by addressing the dependence of the results on the population size. In Fig. (S.2.g), we plot the time average cooperation level in the stationary state, resulted from a simulation in a population of size *N* = 500. Here, the initial condition is a random assignment of the strategies and the preferred games. Here, *g* = 10, *π*_0_ = *c* = 1, and *ν* = 10^−3^. As can be seen, cooperation evolves in smaller population sizes as well. Comparison with Fig. (S.2.c), for the case of a population of size *N* = 20000, reveals that finite size effects favor defection, such that the phase transition line for the defective fixed point to the cyclic orbit shifts towards larger enhancement factors by decreasing the population size. This aspect of the model seems different from many mechanism for the evolution of cooperation, as in the case of many other models on the subject, cooperation level decreases by increasing the population size. This problem is known as the scalable cooperation problem and can be at work to undermine cooperation in large population sizes in many cases (see refs. [29,30] in the main text). In contrast, our analysis suggests competition between public resources, although effective in small population sizes, by solving scalable cooperation problem, can be a strong mechanism to promote cooperation in large populations.

**Fig. S.4.**
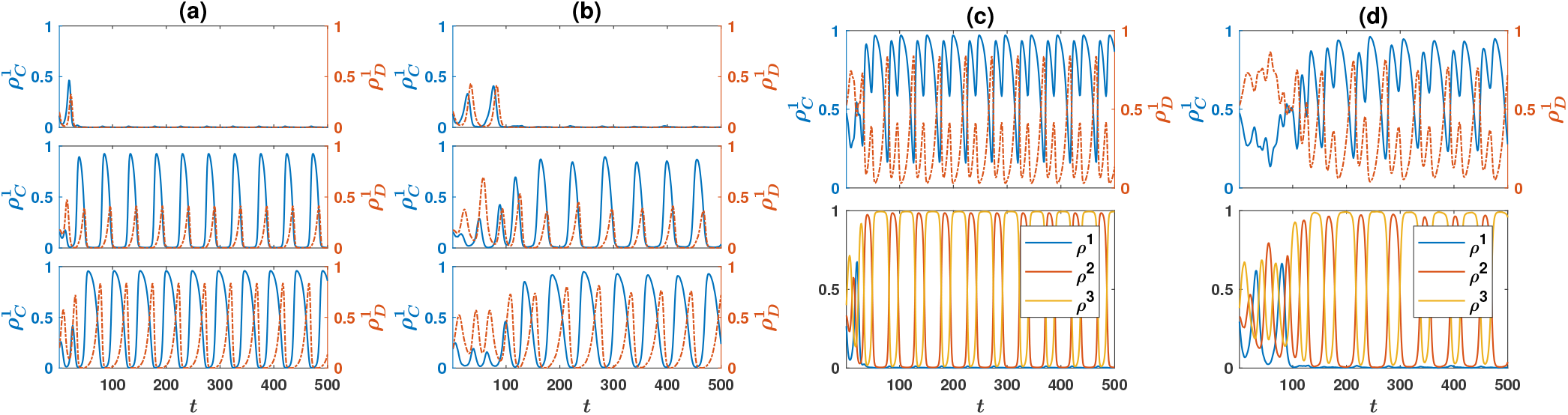
The dynamics of the model for *n* = 3 games. (a) and (b): The density of cooperators 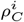, and defectors 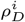 who prefer game *i*, for *i* = 1 to *i* = 3 (from top to bottom). (c) and (d): The total density of cooperators 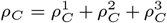 (top), and the density of individuals who prefer different games 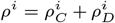, for *i* = 1 to *i* = 3. (a) and (c) result from the replicator dynamics, and (b) and (d) result from a simulation in a population of size *N* = 20000. Here, *g* = 10, *ν* = 0.001, *c* = *π*_0_ = 1, *r*_1_ = 2, *r*_2_ = 2.75, and *r*_3_ = 3.5. The initial condition is a homogeneous initial condition in which the strategies and the preferred games of the individuals are randomly assigned.

To see how the average cooperation level depends on the population size, in Fig. (S.2.h), we plot the average cooperation level over the whole phase diagram (the region defined by 1 ≤ *r*_1_, ≤ *r*_2_ ≤ *g*), as a function of the population size. We note that the initial conditions used in deriving this figure is a homogeneous initial condition in which the strategies and the preferred games of the individuals are randomly assigned. As can be seen, the average cooperation level increases with increasing the population size. As just described, this increment results from a shift in the position of the transition between the defective fixed point and partially cooperative cyclic orbit, to smaller enhancement factors by increasing the population size.

We continue our study of the dependence of the results on the model parameters, by considering the dependence on the group size *g* in Fig. (S.3.a). Here, the contour plot of the cooperation level in the *r*_1_ − *r*_2_ plane, resulted from the numerical solution of the replicator dynamics is presented. Here, we have changed the group size to *g* = 5. As before, we have *ν* = 10^−3^, *c* = *π*_0_ = 1, and the initial condition is a homogeneous initial condition in which the strategies and the preferred games of the individuals are randomly assigned. As expected, the same behavior, observed before for a different group size is observed here: In small enhancement factors, the dynamics settle into a defective fixed point, and a transition to the cyclic orbit in which cooperators survive, occurs as the enhancement factors increase. To see how the cooperation level changes with the group size, we calculate the average cooperation level in the *r*_1_ − *r*_2_ plane, in the interval defined by 1 ≤ *r*_1_*r*_2_ ≤ *g*, and plot this, as a function of group size *g* in Fig. (S.3.c). As can be seen the cooperation level increases with increasing the group size.

To study the dependence of the cooperation level on the mutation rate *ν*, in Fig. (S.3.b), we change the mutation rate to *ν* = 10^−5^, and plot the contour plot of the time average density of the cooperators 〈*ρC*〉_*t*_, as it results from the numerical solutions of the replicator dynamics. Here, we have set *g* = 10, and *c* = *π*_0_ = 1. The initial condition is a homogeneous initial condition in which the density of all the strategies are the same. By comparison with the case of *ν* = 10^−3^ studied before, we note that by decreasing the mutation rate, the phase transition line from the defective fixed point to the cyclic orbit, shifts to smaller enhancement factors (this can be more obviously seen in Figure (1.c) in the main text, where the phase diagram of the model for two different mutation rates are plotted). The shifts in the phase transition lines, result in the enhancement of the cooperation level for smaller mutation rates. This can be seen in Fig. (S.3.d), where the average density of the cooperators in the *r*_1_ − *r*_2_ plane, in the region 1 ≤ *r*_1_, *r*_2_ ≤ *g*, as a function of the mutation rate is plotted. Here, as can be seen, the cooperation level decreases with increasing the mutation rate. However, for very large mutation rates, the cooperation level increases. This is due to the fact that for very large mutation rates, the density of the cooperators maintained in the system due to random mutations is higher than the density of the cooperators maintained in the population as a result of the competition between public resources.

**Fig. S.5.**
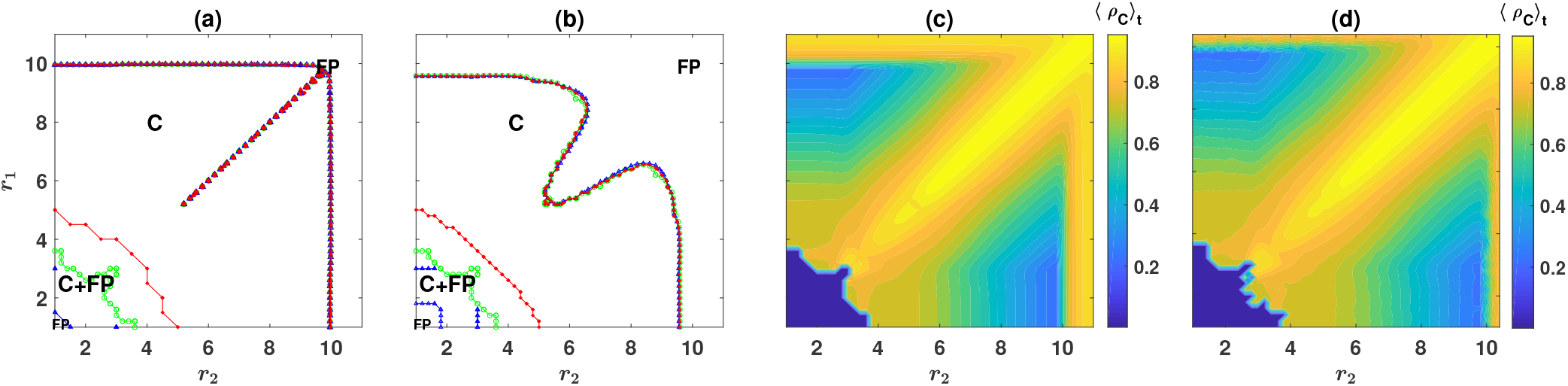
The phase diagram for *n* = 3 games. (a) and (b): The phase diagram of the model for *n* = 3 games, resulted from the replicator dynamics. Here, *g* = 10, *c* = *π*_0_ = 1, *r*_3_ = 3. In (a) *ν* = 10^−5^ and in (b) *ν* = 10^−3^. For small enhancement factors the model shows bistability and can settle in either a defective fixed point, or a cyclic orbit in which cooperators survive. The blue line (marked with small blue triangles) shows the lower boundary of the coexistence region above which the cyclic orbit becomes stable, and the red line (marked with small filled c ircles) s hows t he u pper b oundary o f t he c oexistence r egion, a bove w hich t he d efective fixed po int becomes unstable. The green line (marked with small squares) shows the phase boundary resulted from an unbiased (homogeneous) initial condition. (c) and (d): The contour plots of the time average cooperation level, 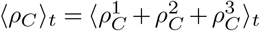, resulting from numerical solutions of the replicator dynamics with *n* = 3 games (c), and a simulation in population of size *N* = 10000. Here, *g* = 10, *c* = *π*_0_ = 1, *r*_3_ = 3, and *ν* = 10^−3^. The replicator equations are solved for *T* = 4000 time steps and the time averages are calculated based on the last 2000 time steps. The simulations are performed for *T* = 5000 time steps, and the time averages are taken over the last 4000 time steps. The initial condition is a homogeneous initial condition in which the strategies and the preferred game of the individuals are randomly assigned.

### Supplementary Note S.6: The case of n = 3 games

As it is made explicit in the main text, the model can be defined in the general context where *n* ≥ 2 PGGs exist and compete to attract individuals. The replicator-mutation dynamics derived here, considers this general case. Here, to shed light on the dynamics of the model in the case that *n* > 2 public resources exist, we consider a case with *n* = 3 public resources, and study the dynamics of the model, using the replicator dynamics and simulations.

We begin by noting that, similarly to the *n* = 2 case, the dynamics with *n* = 3 can settle in a fixed p oint o r a l imit cycle. We begin by studying the cyclic behavior. For this purpose, we plot the density of cooperators 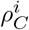, and defectors 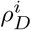, who prefer game *i*, for *i* = 1 to *i* = 3 (from top to bottom), in Fig. (S.4.a) and Fig. (S.4.b). Fig. (S.4.a) shows the result of the replicator dynamics, and Fig. (S.4.b) results from a simulation in a population of size *N* = 10000. The total density of cooperators 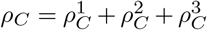, and the total density of defectors 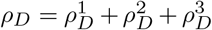 are plotted in the top panels of Fig.(S.4.c) (replicator dynamics) and Fig. (S.4.d) (simulations). The density of the individuals who prefer game *i* is plotted in the bottom panel of Fig. (S.4.c) (replicator dynamics) and Fig. (S.4.d) (simulations). Here, *g* = 10, *ν* = 0.001, *r*_1_ = 2, *r*_2_ = 2.75, and *r*_3_ = 3.5. The initial condition is a homogeneous initial condition in which the strategies and the preferred games of the individuals are randomly assigned.

As can be seen in the figure, the densities of individuals in different public resources shows fluctuations and on average is higher in higher quality resource. This results from the fact that during a cycle a high quality resource attracts and maintains individuals for a longer period of time compared to the low quality resources. However, when a high quality resource becomes occupied with a high density of cooperators, defectors start to grow in that resource. This decreases the profitability of the high quality resource such that eventually cooperators take shelter in a lower quality resource and make it the more profitable resource. We note that, for the values of the enhancement factors used here, the lowest quality resource (game 1, with *r*_1_ = 2) becomes almost obsolete in the stationary state. However, this is not necessarily the case, and it can happen that all the resources attract a significant number of individuals, provided their enhancement factors are very close.

The phase diagram of the model with three games is plotted in Fig. (S.5.a) and Fig. (S.5.b), for two different mutation rates. In Fig. (S.5.a), *ν* = 10^−5^, and in Fig. (S.5.b), *ν* = 10^−3^. Here, we set *g* = 10, and *c* = *π*_0_ = 1, and use the replicator dynamics to drive the phase diagram. The enhancement factor of game *i* = 3 is fixed at *r*_3_ = 3 and the phase diagram is plotted in the *r*_1_ − *r*_2_ plane. For small enhancement factors, the model settles into a defective fixed point where cooperation does not evolve. As the enhancement factor increases, the model becomes bistable and can settle in either the defective fixed point, or a cyclic orbit in which cooperators survive and cyclically dominate the population. The blue line (marked with small blue triangles) shows the lower boundary of the coexistence region above which the cyclic orbit becomes stable. On the other hand, the red line (marked with small filled circles) shows the upper boundary of the coexistence region, above which the defective fixed point becomes unstable and the dynamics settle in the cyclic orbit. In between the two bifurcation lines and in the coexistence region, both the defective fixed point and the limit cycle are stable, and depending on the initial conditions, the dynamics settle into one of these. The phase boundary can be defined as the boundary below which the dynamics settle into the defective fixed point, and above which it settles into the cyclic orbit, starting from an unbiased, homogeneous initial condition. This is indicated in the figure by the green line (marked with small squares). Finally, for large enhancement factors, the model shows a final bifurcation line above which the limit cycle becomes unstable and the dynamics settle in a cooperative fixed point, in which the vast majority of the individuals cooperate.

To see how the cooperation level changes in different regions of the phase diagram, in Fig. (S.5.c) and Fig. (S.5.d), we plot the time average cooperation level, 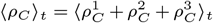. Fig. (S.5.c) represents the result of the replicator dynamics and Fig. (S.5.d) shows the result of simulations. Here, *ν* = 10^−3^, and the same parameter values used in driving the phase diagram are used. The initial condition is a homogeneous initial condition in which the strategies and preferred game of the individuals are randomly assigned. For the replicator dynamics this amounts to an initial condition in which the initial density of all the strategies are equal. The transition between the defective fixed point, with very small level of cooperation, and the cyclic orbit, in which cooperation evolves, is apparent in Fig. (S.5.c) and Fig. (S.5.d). As was the case in the model with *n* = 2 games, the cooperation level is maximized along the diagonal, where the two higher quality games have the same enhancement factors, and it decreases by going away from the diagonal. This shows the importance of the competitiveness of the competing resources in promoting cooperation.

**Fig. S.6.**
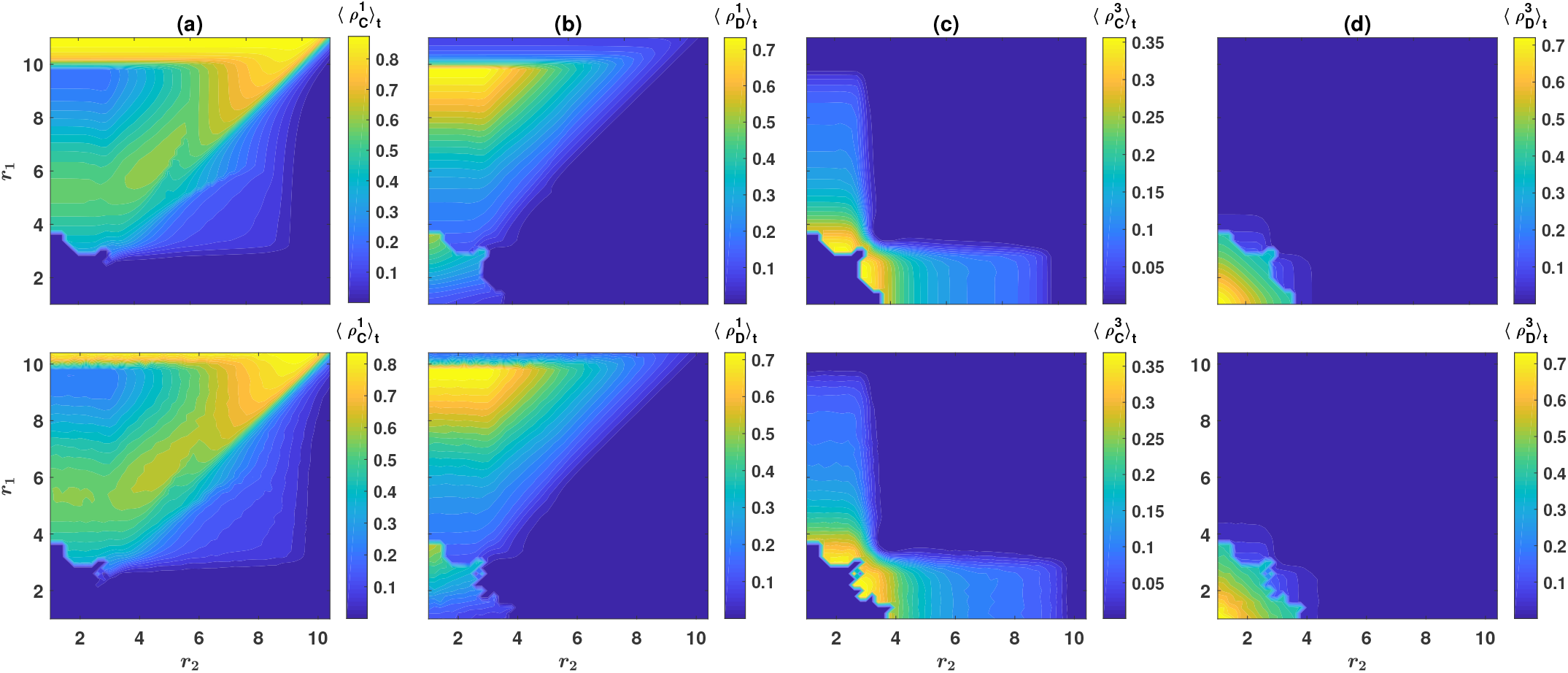
The densities of different strategies for *n* = 3 games. Here, the time average density of cooperators in game 1, 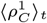, (a), the time average density of defectors in game 1, 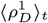, (b), the time average density of cooperators in game 3, 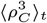, (c), the time average density of defectors in game 3, 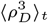, (d), are plotted. The top panels show the results of the replicator dynamics and the bottom panels show the result of a simulation in a population of size *N* = 10000. Here, *g* = 10, *c* = *π*_0_ = 1, *r*_3_ = 3, and *ν* = 10^−3^. The replicator equations are solved for *T* = 4000 time steps and the time averages are calculated based on the last 2000 time steps. The simulations are performed for *T* = 5000 time steps, and the time averages are taken over the last 4000 time steps. The initial condition is a homogeneous initial condition in which the strategies and the preferred game of the individuals are randomly assigned.

The time average density of cooperators in game 1, 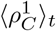, and that of defectors in game 1, 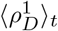 are plotted in, respectively, Fig. (S.6.a) and Fig. (S.6.b). The time average density of cooperators in game 3, 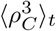, and the time average density of defectors in game 3, 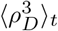 are plotted in, respectively, Fig. (S.6.c) and Fig. (S.6.d). Here, as before, we fix *r*_3_ = 3 and plot the corresponding variables in the *r*_1_ − *r*_2_ plane. The parameter values used here are, as before, *g* = 10, *c* = *π*_0_ = 1, *ν* = 10^−3^ and the initial condition is the random assignment of strategies and the preferred games. The top panels show the result of the replicator dynamics and the bottom panels show the result of a simulation in a population of size *N* = 10000. We note that, due to the symmetry of game 1 and 2, the densities of cooperators and defectors in game 2 result from that in game 1, in Fig. (S.6), by reflection with respect to the diagonal. As it can be seen, while for small enhancement factors cooperation does not evolve, as the enhancement factors increase, a phase transition to a phase where cooperation does evolve occurs. Interestingly, in the region where both *r*_1_ and *r*_2_ are larger than *r*_3_ = 3, that is when game 3 is the most inferior resource, it becomes almost obsolete: very few individuals prefer game 3, and the dynamics is determined by interaction of the two higher quality resources. However, although game 3 does not attract individuals significantly, the mere possibility of playing a third game have a positive effect on the cooperation level. This can be seen by comparison with the case of *n* = 2 games. On the other hand, in the region where either *r*_1_ or *r*_2_ are larger than *r*_3_ = 3, that is when game 3 is the second high quality resource, it always attracts cooperators, but not defectors (compare Fig. (S.6.c) and Fig. (S.6.d)). This results from the fact that defectors prefer the highest quality resource. This can be seen by looking at the region *r*_1_ *> r*_2_ in Fig. (S.6.a) and Fig. (S.6.b), where respectively 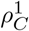 and 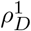 are plotted. As can be seen in this region, both cooperators and defectors prefer game 1, the highest quality resource. This interesting observation hints at the mechanism by which competition between resources leads to the evolution of cooperation: When individuals can choose between different resources, defectors tend to prefer the highest quality resource, and are unwilling to choose the other less quality resources. This makes a second low quality resource a shelter for cooperators where they can work cooperatively and grow safely, when the density of defectors increases in the population. The lowest quality resource on the other hand, remains almost obsolete (as long as its quality is sufficiently lower than the other two resources). This is due to the fact that the second quality resource, having a low fraction of defectors, is already a good enough shelter for cooperators and they do not need to switch to the lowest quality resource.

So far we have seen that cooperators can survive in the population by switching to the second high quality resource when the density of defectors increase in the population. However, when the enhancement factor of the second quality resource is much lower than that of the highest quality resource, it becomes more difficult for cooperators to out-compete defectors by sheltering in the low quality resource and making it the most profitable one. This leads to the fact that the cooperation level is the highest when the enhancement factors of the two highest quality resources are similar, and it decreases by increasing the difference between the enhancement factors of the two high quality resources (see Fig. (S.5.c) and Fig. (S.5.d)). This leads to the fact that, for a fixed *r*_3_ (smaller than *g*), the cooperation level in the system, as well as the cooperation level in game 1 is maximized when the enhancement factors of the two highest quality resources are the same. Increasing *r*_1_ or *r*_2_ beyond this value, has a detrimental effect on the level of cooperation and increases the density of defectors in the highest quality resource (see Fig. (S.6.b)), and thus, in the population.

## Notes

### Competing Interest Statement

The authors have declared no competing interest.

